# Scalable sequence-informed embedding of single-cell ATAC-seq data with CellSpace

**DOI:** 10.1101/2022.05.02.490310

**Authors:** Zakieh Tayyebi, Allison R. Pine, Christina S. Leslie

## Abstract

Standard scATAC-seq analysis pipelines represent cells as sparse numeric vectors relative to an atlas of peaks or genomic tiles and consequently ignore genomic sequence information at accessible loci. We present CellSpace, an efficient and scalable sequence-informed embedding algorithm for scATAC-seq that learns a mapping of DNA k-mers and cells to the same space. CellSpace captures meaningful latent structure in scATAC-seq datasets, including cell subpopulations and developmental hierarchies, and scores the activity of transcription factors in single cells based on proximity to binding motifs embedded in the same space. Importantly, CellSpace implicitly mitigates batch effects arising from multiple samples, donors, or assays, even when individual datasets are processed relative to different peak atlases. Thus, CellSpace provides a powerful tool for integrating and interpreting large-scale scATAC-seq compendia.

## Main

Typical computational strategies to discover latent structure in scATAC-seq datasets mimic scRNA-seq workflows. First, scATAC-seq data is summarized as a sparse numeric cell-by-event matrix, where the events correspond either to an atlas of chromatin accessible peaks or to highly variable genomic tiles^1, 2^, analogous to the cell-by-gene matrix in scRNA-seq analysis. The cell-by-event matrix can be binarized (1 indicating that the event was accessible in a cell, 0 that the event was inaccessible or not captured) or contain counts. Then normalization followed by a standard dimensionality reduction method such as latent semantic indexing (LSI) yields a lower-dimensional representation for cells, allowing construction of a nearest neighbor (NN) graph on cells based on similarity in the lower-dimensional space and application of existing graph-based clustering and embedding algorithms from the scRNA-seq toolkit. However, due to its high dimensionality and sparsity, dimensionality reduction and embedding of scATAC-seq is challenging and prone to complex batch effects. Other analysis methods, in particular those that seek to integrate scATAC-seq and scRNA-seq datasets, summarize single-cell chromatin accessibility profiles at the gene promoter or locus level to generate scRNA-seq-like data^3^. This approach loses the representational richness of scATAC-seq, namely the accessibility of cell type specific enhancers and the transcription factor (TF) binding motifs they contain.

Rather than use these scRNA-seq-inspired strategies, we seek to move beyond the cell-by-event count matrix representation of scATAC-seq and exploit the genomic DNA sequences underlying accessible peaks/tiles, since sequence features like TF binding motifs are informative of developmental state and cell identity and therefore should help reveal biologically meaningful latent structure. Importantly, we will incorporate genomic sequence information in the latent structure discovery step of scATAC-seq analysis, rather than in a post hoc analysis step. To date, very few approaches have attempted sequence-informed structure discovery or embedding of scATAC-seq. Early work used chromVAR^4^ to represent each cell as a vector of accessibility scores relative to a fixed library of known TF motifs^5^. This approach can indeed group cells by their cell type identity but introduces bias through a priori motif choice and therefore may miss latent structure in the data. In addition, TF motif accessibility scores can capture technical differences between samples and hence preserve batch effects. Recently, scBasset^6^ used a multi-task neural network approach to learn a sequence model for chromatin accessible peaks that passes through a low-dimensional bottleneck layer, together with cell-specific model vectors that predict whether a peak—given its bottleneck representation—will be accessible in the cell. This approach yields a low dimensional representation of cells via the model vectors and can assign TF accessibility scores to cells by passing a sequence with a planted motif through the model. However, scBasset requires training of a large neural network model where the number of tasks equals the number of cells and likely will require further optimizations to scale to large datasets. Finally, a recent method called SIMBA uses a graph embedding approach for scRNA-seq, scATAC-seq, and multiome data^7^, where cells, genes, peaks, *k*-mers, and TF motifs are vertices in the graph for a dataset, and edges connect entities (like peaks) that relate to other entities (like cells). Notably for the application of this method to scATAC-seq, the TF motifs must be specified prior to training in order to define the graph and therefore will impact the learned embedding. Moreover, the cell-by-peak matrix is explicitly encoded in the graph, potentially tying the approach to the inherent sparsity and batch effect issues of the cell-by-peak representation.

Here we present CellSpace, an efficient and scalable *k*-mer based embedding algorithm for scATAC-seq. CellSpace employs a latent embedding algorithm from natural language processing called StarSpace^8^, similar to the strategy we used in the BindSpace model for embedding TF SELEX-seq data to represent subtle binding preferences of TFs^9^. CellSpace learns a joint embedding of *k*-mers and cells so that cells will be embedded close to each other in the latent space not simply due to shared accessible events but based on the shared DNA sequence content of their accessible events. Notably, CellSpace avoids explicitly embedding peaks/tiles, and therefore does not encode the cell-by-event matrix. Single cell TF motif activities can be readily computed in CellSpace’s latent space; the selection of TF motifs is not required ahead of time and does not influence training. Importantly, thanks to key representational and training choices, we show that CellSpace’s sequence-aware embedding has powerful intrinsic batch mitigating properties, allowing discovery of latent structure to enable trajectory analysis and cluster discovery in single cell chromatin accessibility datasets across multiple samples and assays, even when the individual datasets are processed independently.

## Results

CellSpace trains on scATAC-seq data to learn an embedding of DNA *k*-mers and cells into a common latent space (**Fig. 1**, **Methods**). To generate training examples, CellSpace samples genomic sequences of fixed length from accessible events (peak atlas or genomic tiles) and uses the set of cells in which an event is present as positive labels for the sampled input sequence (**Fig. 1a**). This process produces (LHS, RHS) training pairs where the left-hand side is a bag of *k*-mers representation of the sampled sequence, and the right-hand side is a cell in which the event is accessible. During training, CellSpace updates the embedding vectors of *k*-mers and cells to push the induced embedding representation of the LHS sequence towards the embedding of the ‘positive’ cell on the RHS and away from cells sampled from all ‘negative’ cells (**Fig. 1b**). Here, a *K*-negative sampling strategy^10^, where *K* negative cells are sampled at random, is used to improve training time by updating only some of the weights at each optimization step. This technique is often used for data with orders of magnitudes more negative observations compared to positive ones, which is the case for the accessibility vector of most events. Moreover, sampling from the negative cells, instead of using all cells for which the event is labeled 0, reduces the effect of false negatives caused by scATAC-seq sparsity. Importantly, CellSpace uses *N*-grams in the bag of *k*-mers representation to extract context from the data, so that considering the proximity of *k*-mers improves the embedding (**Fig. 1b**).

**Figure 1.**
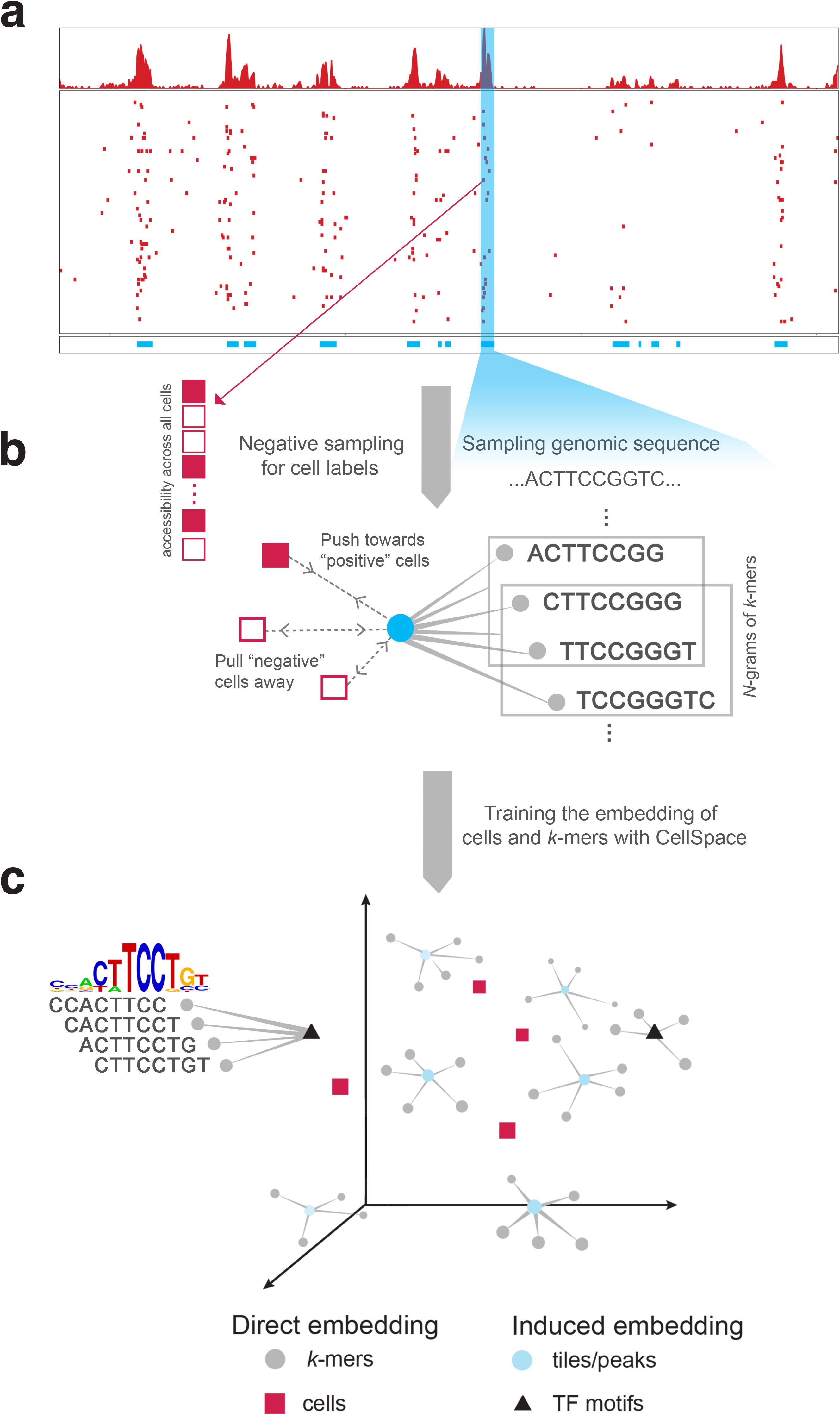
CellSpace learns a sequence-informed embedding of cells from scATAC-seq. Overview of the CellSpace algorithm. **a.** CellSpace samples sequences from accessible events (peaks or tiles) to generate training examples, each consisting of an ordered list of overlapping *k*-mers from the sampled sequence, a positive cell (where the event is open), and a sample of negative cells (where the event in closed). **b.** CellSpace learns an embedding of *k*-mers and cells into the same latent space. For each training example, the embeddings of the corresponding *k*-mers and cells are updated to pull the induced sequence embedding towards the positive cell and away from the negative cells in the latent space; learning contextual information, represented by *N*-grams of nearby *k*-mers, improves the embedding. **c.** Once the embedding of cells and *k*-mers is trained, TF motifs can be mapped to the latent space, allowing cells to be scored for TF activities based on TF-cell similarities.

Accessible events (peaks/tiles) are not explicitly embedded; an induced representation of an event can be computed from the embedding of its DNA sequence *k*-mers. By not directly embedding peaks—and by updating the cell embedding based on the *k*-mer content of accessible regions rather than the identity of these regions—CellSpace appears to be less influenced by the choice of preprocessing pipeline for peak calling or variable tile selection and by technical differences between batches and even assay variants. Finally, any TF motif can be embedded in the latent space based on the embedding of the constituent *k*-mers from its consensus sequence (**Fig. 1c**). Notably, this means that the set of TF motifs to be examined is not required at training time, and the embedding is not affected by a specific set of known motifs. Similarity between a TF motif embedding and cell embedding in the latent space produces a TF activity score, and these motif scores are useful in characterizing cell subpopulations. Finally, similarity of cells in the latent space can be used to produce a NN graph for clustering, visualization with UMAP^11^, and other downstream analyses (**Fig. 1c**).

We first tested our approach on a smaller scATAC-seq dataset profiling CD34+ hematopoietic stem and progenitor cell (HSPC) populations from multiple human donors^5^, where ground truth cell types based on fluorescence activated cell sorting (FACS) are available. After filtering steps, doublet removal, and variable tile (500 bp genomic bin) selection using ArchR, we retained 2,154 cells for CellSpace embedding using 50K variable tiles, sampling 150 bp sequences with 3-grams of 8-mers. CellSpace obtained a biologically meaningful embedding of the hematopoietic differentiation hierarchy as visualized by UMAP (**Fig. 2a**), where hematopoietic stem cells (HSC) and multipotent progenitors (MPP) diverge into two main erythroid and lymphoid branches, with common myeloid progenitors (CMP) giving rise to megakaryocyte-erythrocyte progenitors (MEP) along one branch and lymphoid-primed multipotent progenitors (LMPP) giving rise to common lymphoid progenitors (CLP) along the other. The granulocyte-monocyte progenitors (GMP) branch off both from LMPP and CMP populations, consistent with current models of hematopoiesis (**Fig. 2b**). Trajectory analysis with Palantir^12^, using a cell from the HSC population as the origin, recovers six termini that include the most differentiated cell types in the hierarchy that are represented in the dataset: CLP, plasmacytoid dendritic cells (pDC), MEP, an endpoint within the GMP population and a GMP-adjacent population labeled as “unknown” in the original study, and monocytes (**Fig. 2c**, **Supplementary Fig. 1a**). We also embedded motifs for TFs important in hematopoietic differentiation using CellSpace, as shown in the CellSpace UMAP embedding (**Fig. 2a**). We note that inclusion of motif ‘vertices’ in the NN graph very slightly changes the 2-dimensional UMAP visualization. The location of motifs in the UMAP gives an intuition for why CellSpace recovers the developmental hierarchy correctly, as cell-type-specific TFs are embedded close to the cells where they are active: for example, the HOXA9 motif is embedded near the HSC population, GATA1 near MEP, CEBPB near GMP, PAX5 near CLP, and IRF1 near pDC. Moreover, TFs active in multiple cell types end up in between them: for example, the ESRRA motif is close to GMP and pDC populations.

**Figure 2.**
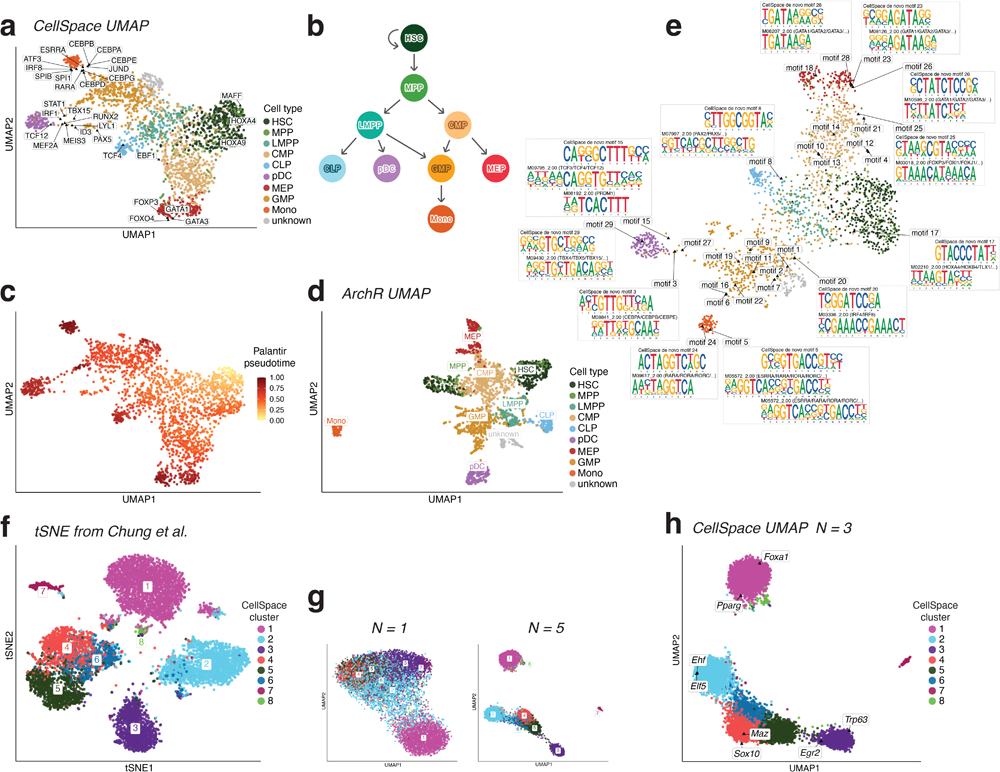
CellSpace recovers latent structure and developmental hierarchies. **a.** UMAP of CellSpace embedding for 2,154 cells from human HSPC scATAC-seq data annotated by FACS-sorted cell types. Embedding of key hematopoietic TF motifs also shown. **b.** Current model of hematopoietic differentiation, with cell labels and colors as in (**a**). **c.** Palantir pseudotime analysis using CellSpace embedding, with an HSC starting point, identifies differentiation termini corresponding to CLP, pDC, GMP, MEP, and monocyte fates. **d.** UMAP of iterative LSI embedding based on cell-by-tile matrix using ArchR splits HSC, MPP, and MEP populations into two clusters due to batch effects. **e.** UMAP of cells and *de novo* motifs discovered based on the same trained CellSpace embedding as DNA 10-mers that are frequent nearest-neighbors of each cluster’s cells are identified and clustered by sequence content; 10-mer clusters are aligned and converted to PWMs. **f.** Standard t-SNE from LSI dimensionality reduction of the cell-by-peak matrix from the mammary epithelial scATAC-seq dataset. Cells are annotated using CellSpace clusters (*N* = 3), and comparison with the original study was used to associate these clusters with cell types. **g.** UMAP of CellSpace embedding for 7,846 cells from murine fetal and adult mammary epithelial scATAC-seq data shows the impact of *N*-gram parameter for *N* = 1, 5. **h.** CellSpace with default *N* = 3 accurately captures developmental relationships between cell types. Key TF motifs in epithelial differentiation also shown in the *N* = 3 CellSpace embedding.

Strikingly, CellSpace mitigates batch effects in this dataset, with cells from multiple donors well mixed and in particular with HSC/MPP populations from three donors clustering together (**Supplementary Fig. 1b**). Indeed, Seurat SNN clustering based on the CellSpace embedding largely recovered the known cell type labels, with the earliest stem and progenitor populations HSC and MPP grouping in one cluster (**Supplementary Fig. 1b**). By contrast, iterative LSI (itLSI) dimensionality reduction using ArchR separated the HSC/MPP populations into two separate clusters, segregating one of the donors from the other and obscuring the overall hierarchy (**Fig. 2d**, **Supplementary Fig. 1c**). Similarly, scBasset reported a strong donor batch effect in their embedding of this dataset, and a modification of the model to explicitly account for batch was required to overcome this issue^6^.

We also asked whether we could learn TF motifs *de novo* from the CellSpace embedding—rather than embedding known TF motifs—since such a procedure could in principle allow the discovery of novel motifs. To do this, we used the trained CellSpace embedding on the Buenrostro *et al.* dataset to find the induced embedding of all DNA 10-mers, and for each cell cluster (**Supplementary Fig. 1b**), compiled the 10-mers that are frequently among the nearest-neighbors of cells in the cluster (**Methods**). Next, we clustered these 10-mers based on their sequence composition, aligned the 10-mers in each cluster, and computed a position weight matrix (PWM) from each alignment, yielding 29 *de novo* motifs (**Supplementary Fig. 1d, Methods**), which we visualized in the CellSpace embedding using UMAP (**Fig. 2e**). After comparing this *de novo* motif discovery procedure to CIS-BP^13^ motifs, we found that we retrieved motifs similar to many of the relevant hematopoietic TF motifs (**Fig. 2e**), suggesting the potential for learning meaningful novel motifs in systems where important factors are unknown.

To quantify the extent to which CellSpace implicitly corrects batch effects while preserving biological heterogeneity, and to compare to other recent scATAC-seq embedding methods, we assessed the batch effect using published metrics (kBET, bASW, GC)^14^ as well as a mutual information based metric (bNMI) and also evaluated clustering quality metrics (HG, ARI, NMI, ASW)^14, 26^ (**Methods**). Successful batch mitigation or explicit batch correction should yield good scores for the batch correction metrics without sacrificing biological complexity, including the biological coherence of clusters as assessed by the clustering metrics. To statistically assess differences in performance, we used aggregated scores— producing a single metric for batch, a single metric for clustering/biological complexity, and a single overall metric—and performed a bootstrapping analysis to report 95% confidence intervals and FDR-adjusted *p*-values for pairwise comparisons between algorithms (**Supplementary Fig. 2a-e, Supplementary Tables 2,3, Methods**). The “unknown” cell type was included in embedding but excluded from evaluations. We assessed CellSpace embeddings based on variable genomic tiles and on variable peaks and compared to a wide range of existing methods: ArchR’s itLSI using variable tiles; standard LSI using peaks; scBasset; SIMBA using either peaks alone or peaks, *k*-mers, and TF motifs in the graph embedding; PeakVI^15^, a variational autoencoder embedding of the cell-by-peak matrix; and chromVAR using motifs or *k*-mers. For methods that provide an explicit batch correction approach (scBasset, SIMBA, PeakVI), we ran both without the batch covariate, to assess implicit batch mitigation, and with the batch covariate, to assess explicit batch correction. For LSI-based embedding methods, we also evaluated metrics after batch correction with Harmony^16^, a widely-used single-cell integration method.

We found that CellSpace (variable tiles) significantly outperforms scBasset (with and without batch correction, adjusted *p* < 0.05 and 0.01, respectively), all variants of SIMBA (adjusted *p* < 0.05 to 0.01), PeakVI (with and without batch correction, adjusted *p* < 0.05 and 0.01, respectively), both variants of chromVAR (adjusted *p* < 0.01), and LSI (peaks) without batch correction (adjusted *p* < 0.05) (**Supplementary Fig. 2d**). Based on bootstrap analysis, CellSpace (variable tiles) is significantly better than ArchR itLSI (variable tiles) in terms of batch correction (adjusted *p* < 0.01), but there is no significant difference in terms of the biological complexity score and overall score between these methods. CellSpace, which uses no knowledge of batch covariates, performs comparably on this small data to Harmony batch correction applied to ArchR itLSI (variable tiles) or LSI (peaks). CellSpace (variable tiles) has a higher average overall score than all other methods, even in cases where the adjusted *p*-value is higher than 0.05. Note that all variants of LSI (applied to peaks or variable tiles and with or without batch correction) are not sequence-informed embeddings and do not provide batch-corrected TF motif scores, whereas CellSpace provides TF motif scores that do not require an explicit batch correction.

We also show individual batch metrics by cell type (**Supplementary Fig. 2e**), for metrics that are averaged over all cell types, which we provide for intuition without confidence intervals. Of the competing methods to CellSpace (variable tiles), only the approaches with explicit batch correction improve the batch metrics for the HSC and MPP cell types, which are most affected by donor batch; in some cases (e.g., ArchR itLSI + Harmony, batch-corrected SIMBA), improvement for HSC and MPP seems to come at the cost of poorer performance on MEP, which is another cell type with a donor batch effect. Overall, CellSpace (variable tiles) either ties or significantly outperforms all competing methods on this dataset, including methods with explicit batch correction using the batch (‘donor’) covariate, and notably outperforms sequence-informed methods that provide TF motif scores.

We found that the use of *N*-grams in the CellSpace algorithm was often important for recovering well-defined latent structure in the embedding. To illustrate this effect, we applied CellSpace to a second published scATAC-seq dataset profiling 7,846 murine fetal and adult mammary epithelial cells using the peak atlas provided by the authors^17^. We first reproduced the t-SNE visualization of the dataset from the original study using standard processing of the cell-by-peak matrix to identify the reported cell types: adult luminal progenitor (LP), adult mature luminal (ML), adult basal, LP-like fetal, ML-like fetal, and basal-like fetal (**Fig. 2f**). Next, we ran CellSpace with different choices of the *N*-gram hyperparameter, sampling *L* = 300 bp sequences (instead of the default *L* = 150 bp) due to the larger size of the peaks (1000 bp) and plotted UMAP visualizations (**Fig. 2g,h**). We found that *N* = 1 (**Fig. 2g**, simple bag of 8-mers) yielded a diffuse embedding, while *N* = 3 (**Fig. 2h**, default) clarified the population structure and identified correct developmental relationships between fetal and adult cell types. The larger value *N* = 5 (**Fig. 2g**) began to pull cell populations further apart in the embedding, although clustering and developmental relationships were still correct. Canonical luminal (Foxa1, Pparg) and basal (Trp63, Egr2) TFs were correctly associated with cell populations via the CellSpace motif embeddings (*N* = 3, **Fig. 2h**).

Beyond visualizing TF motifs in the CellSpace embedding with UMAP, we can compute single cell TF activity scores by computing the similarity between TF motif and cell embeddings in the latent space (**Methods**). To systematically assess CellSpace’s motif scoring, we next analyzed a recent multiome dataset profiling the human cortex containing 8,981 cells with both scRNA-seq and scATAC-seq readouts^18^. Running CellSpace with default parameters (*L* = 150 bp, *N* = 3) on the provided scATAC-seq cell-by-peak matrix readily captured major developmental relationships between cell types based on reported cluster annotations, with glutamatergic neuron (GluN) clusters grouping apart from inhibitory neuron (IN) clusters in the UMAP (**Fig. 3a**). We also compared to the UMAP visualization of the scBasset embedding of the same scATAC-seq dataset. We found that scBasset converged before 45 epochs (before the default 1000 epochs, **Supplementary Fig. 3a**) and trained efficiently when using specialized large-memory GPUs (**Supplementary Table 1**). Notably, scBasset applies stringent filtering to the training data, effectively decreasing the number of peak training examples by an order of magnitude. scBasset found a topologically similar embedding to CellSpace, but unlike CellSpace and the standard LSI embedding, it failed to separate inhibitory neuron cluster IN3 from the glutamatergic neurons (GluN) (**Fig. 3a**).

**Figure 3.**
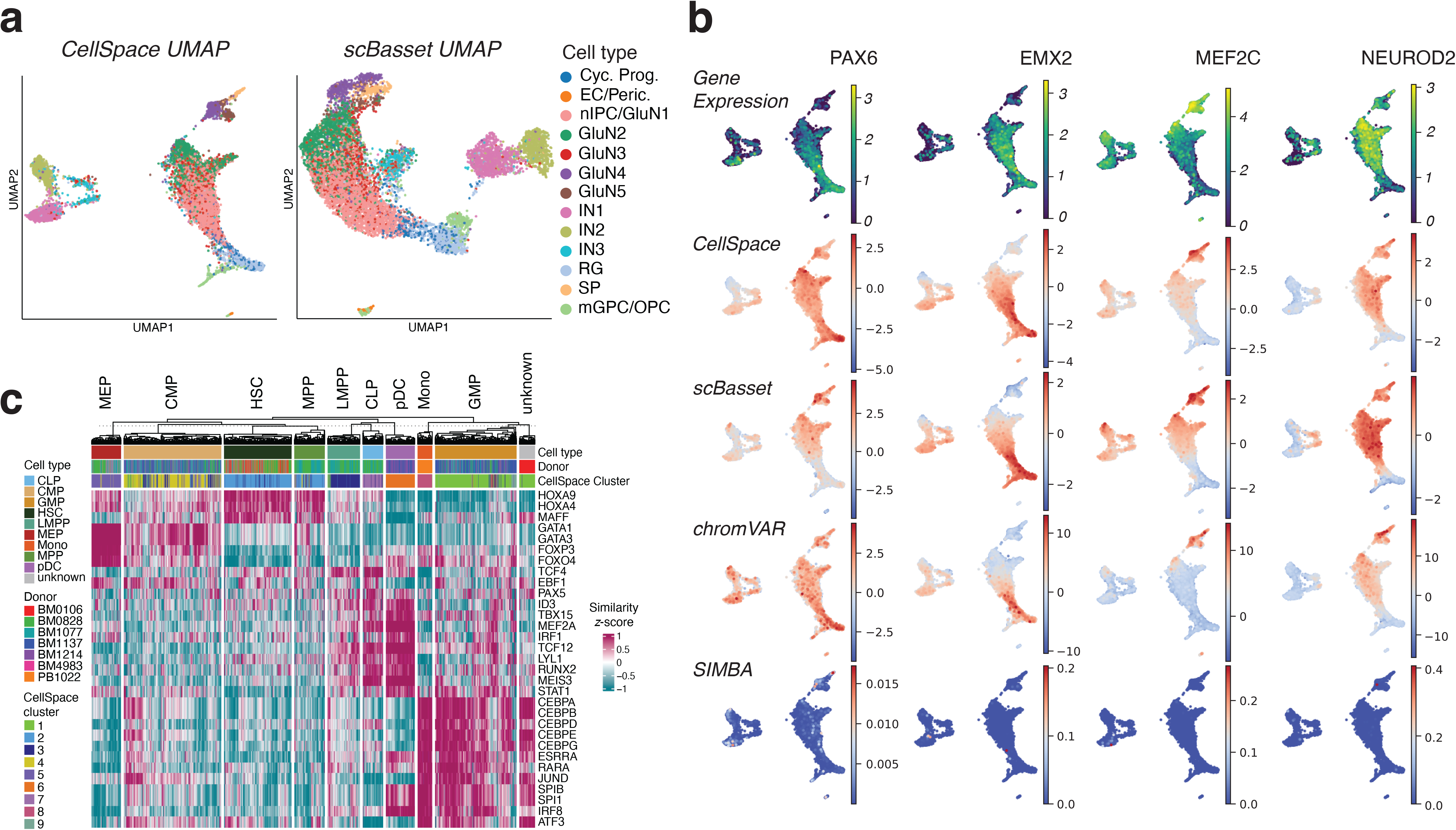
Single cell motif scoring using CellSpace accurately maps TF activities. **a.** CellSpace and scBasset embeddings of the scATAC-seq readout of a human cortex multiome dataset with 8,981 cells. **b.** Rows show for the TFs PAX6, EMX2, MEF2C, and NEUROD2, visualized on the CellSpace embedding, gene expression for the TFs; CellSpace motif scores; scBasset motif scores; chromVAR motif deviation scores; SIMBA motif scores. **c.** CellSpace TF motif scoring for hematopoietic dataset, shown as heatmap (cell types and clusters visualized in Fig. 2a, Supplementary Fig. 1b).

Moreover, when we compared the TF motif scores provided by other sequence-informed embedding methods, CellSpace motifs scores for key TFs correlated better with expression of the corresponding factors from the scRNA-seq readout, as exemplified by the corresponding UMAP visualizations (**Fig. 3b**). For example, CellSpace correctly captures that the strongest PAX6 activity is in the radial glia (RG) population, while scBasset associated PAX6 to cell populations where it is not expressed. For EMX2 and MEF2C, CellSpace better captures the overall landscape of TF activity, while scBasset seems to overestimate activity in IN subpopulations. In some cases, such as NEUROD2, both methods correctly map the region of TF activity as validated by expression. For an overall comparison, we computed the correlation between gene expression (from the RNA-seq readout) and TF motif scores (from the ATAC-seq readout) from each method, and considered the set of important neurodevelopmental TFs that were identified by the original authors^18^, restricting to those whose motifs passed scBasset’s default filtering steps (**Methods**). **Supplementary Fig. 3b** shows the correlation values for each method (CellSpace on the y axis and other methods on x axes). CellSpace’s motif scores appear to outperform scBasset’s scores on the important neurodevelopmental factors; in particular, CellSpace TF motifs scores yield positive correlation with expression for almost all these factors (17/19, upper half plane of scatterplots), in contrast with scBasset (14/19 with positive correlation, right half plane of scatterplots). CellSpace’s TF activities had similar performance as chromVAR TF motif deviations in terms of correlation with expression (**Fig. 3b**, **Supplementary Fig. 3b**). To confirm that CellSpace’s improvement over scBasset in motif scoring was not simply due to use of a larger training set of peaks, we reran scBasset on the full peak atlas, which required over twice as long to train and an order of magnitude more memory than CellSpace (**Supplementary Table 1**); this embedding more successfully separated cluster IN3 from the GluN clusters, but the incorrect association of PAX6 to non-RG populations and the over-scoring of EMX2 in the IN1 population persisted (data not shown). Finally, we trained a SIMBA embedding on the peak atlas using *k*-mers and TF motifs. SIMBA had a significantly higher memory usage than CellSpace, but trained faster using peaks associated with top PCs (**Supplementary Table 1**). Moreover, we found that SIMBA motif scores did not provide meaningful per-cell motif activities but rather yielded mostly zero scores across the atlas (**Fig. 3b**); consequently, the correlation of SIMBA TF motif scores with TF expression were mostly close to zero (**Supplementary Fig. 3b**), although SIMBA scores could be used to find an association with cell type via ranking (**Supplementary Fig. 3c**).

To both qualitatively and quantitatively compare across scATAC-seq embedding approaches, we produced UMAP visualizations, clustered cells, computed performance scores for CellSpace, LSI, scBasset, SIMBA (peaks only), SIMBA (peaks+*k*mers+motifs), PeakVI, and chromVAR (**Supplementary Fig. 3d,e, Supplementary Tables 2,4, Methods**), and performed a bootstrapping analysis to report 95% confidence intervals for the overall biological complexity score and FDR-adjusted *p*-values for pairwise comparisons as before. On this dataset, CellSpace (peaks) significantly outperforms LSI (adjusted *p* < 0.01), SIMBA (peaks) (adjusted *p* < 0.05), PeakVI (adjusted *p* < 0.01), and chromVAR (adjusted *p* < 0.01). While the 95% confidence interval of CellSpace is higher than that of SIMBA (peaks+kmers+motifs) and almost non-overlapping with the top of the scBasset confidence interval, the performance difference between these methods is not statistically significant (adjusted *p* = 0.087 for both). Again, CellSpace ties or significantly outperforms all competing methods on the human cortex dataset.

Returning to the hematopoietic dataset previously described (**Fig. 2a**), we can similarly compute motif scores for key developmental TFs in the blood system (**Fig. 3c**). This analysis retrieved the correct association between TFs and HSPC populations, including GATA1 with MEP cells, ID3 with CLP and pDC cells, and CEBPB with GMP cells. Interestingly, a subset of cells in the CMP population that are placed by CellSpace in cluster 1—predominantly made up of GMP cells—indeed have high CEBPB scores, suggesting progression towards the GMP cell state. Motif scoring for the mammary epithelial dataset (**Fig. 2h**) similarly identified correct activities of key luminal and basal TFs in fetal and adult cell populations (**Supplementary Fig. 3f**).

Next, to assess CellSpace’s scalability and batch-mitigating capabilities, we ran the model on several large-scale multi-sample datasets with challenging batch effects. First we turned to a larger human hematopoietic dataset comprised of 61,806 cells collected from bone marrow and peripheral blood from 12 healthy donors^19^ together with 2,706 cells from the smaller HSPC dataset^5^. The cell-by-peak matrix was originally processed in multiple steps, with LSI dimensionality reduction followed by a batch correction procedure, variable peak selection, then re-computing LSI^19^. Cells were then clustered into 31 clusters in this final lower dimensional space; the resulting UMAP with major clusters is reproduced here (**Fig. 4a**). While developmental relationships can be inferred from this embedding, there also appears to be artifactual structure from residual batch effects and noise.

**Figure 4.**
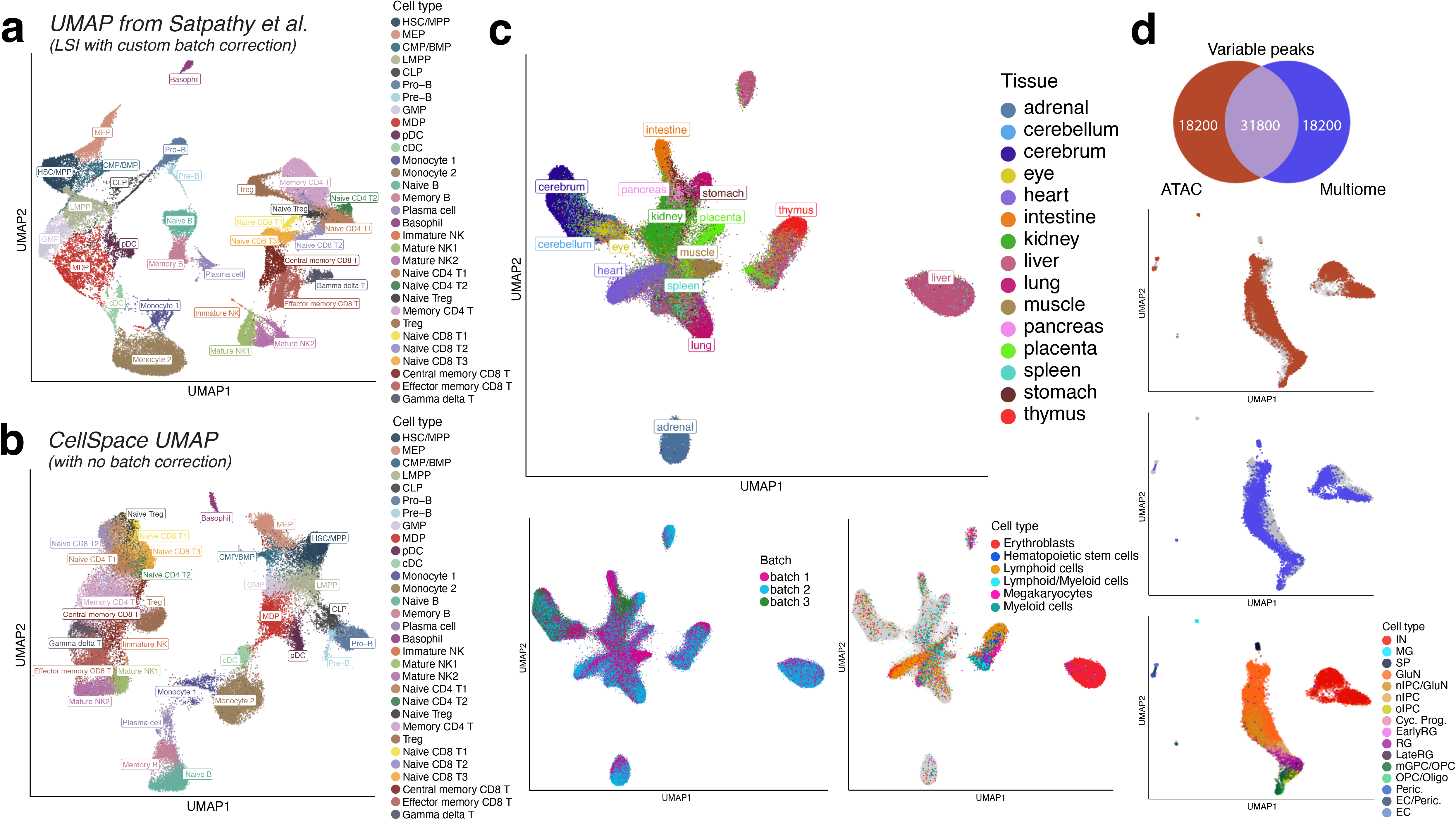
CellSpace’s embedding implicitly mitigates donor-and assay-specific batch effects in large-scale scATAC-seq datasets. **a.** UMAP of LSI dimensionality reduction with custom batch correction from original study of a large-scale multi-donor human hematopoietic dataset with 63,882 cells, annotated with major reported clusters. **b**. CellSpace embedding of large hematopoietic dataset without any custom preprocessing recovers hematopoietic developmental hierarchy. **c.** UMAPs for CellSpace embedding of ∼720K cell human fetal tissue atlas data set from Domcke *et al.* labeled by tissue, by batch, and by blood cell types across multiple tissues. **d.** CellSpace applied to human cortex chromatin accessibility data using two assays, multiome with 8,981 cells and scATAC-seq with 12,675 cells, processed with respect to their own peak atlases. Venn diagram for the top 50K most variable peaks from each assay, with 31,800 peaks in each atlas having non-zero overlap with the other atlas. UMAPs show good mixing of datasets and cell type annotations in the CellSpace joint embedding.

We asked whether CellSpace’s *k*-mer based embedding could overcome batch effects and find latent structure without multiple custom preprocessing steps. We therefore ran CellSpace on this large-scale ∼63K cell dataset using the cell-by-peak matrix for the top 50K variable peaks and with default parameters, except for increasing the embedding dimensions and the number of epochs due to the large number of cells from various populations (**Methods**). Here we exploit the fact that CellSpace is memory-efficient even for large-scale datasets (**Supplementary Table 1**) since random training examples are generated at every step of optimization and only the sparse count matrix and its corresponding genomic sequences are indexed and stored in memory (**Methods**). This is true even if all peaks from this dataset are used instead. A UMAP visualization of the resulting embedding shows that CellSpace faithfully captured the hematopoietic developmental hierarchy within the HSPC compartment and correctly linked progenitor populations to more mature blood cell types (**Fig. 4b**). For example, the monocyte-dendritic progenitor (MDP) population was embedded near to monocytes and conventional dendritic cells (cDC), while common lymphoid progenitor cells (CLP) displayed a differentiation trajectory towards pro-B and pre-B cells. While this dataset contained both the older and newer scATAC-seq profiles and multiple human donors, we found that batches/donors were well mixed in the embedding (**Supplementary Fig. 4a**). Due to the diversity of this dataset, with both bone marrow derived HSPC samples and more differentiated cell types from peripheral blood, we were able to obtain more resolution by retraining the CellSpace embedding on specific compartments. For example, a UMAP visualization of the CellSpace embedding of natural killer (NK) and T cells from this dataset reveals detailed relationships between cell types (**Supplementary Fig. 4b**).

As another example, we applied CellSpace to a scATAC-seq dataset profiling the tumor immune microenvironment (TME) in basal cell carcinoma (BCC) biopsies from 7 patients^19^, comprising 37,818 cells. Although the authors reported a detectable batch effect that confounded further analyses and required attenuation^19^, we ran CellSpace directly on 50K variable peaks and recovered the identified T cell types as well as other lymphoid, myeloid, endothelial, and fibroblast populations that were well-mixed over donors (**Supplementary Fig. 4c**). As has been described in scRNA-seq analyses of tumor samples, the tumor cells from different patients retained more distinct identities in the embedding. These examples show that CellSpace readily scales to large and complex scATAC-seq sets to mitigate batch effects and discover meaningful global latent structure.

We again assessed CellSpace’s batch mitigation properties by comparing biological complexity, batch correction, and overall metrics against both sequence-informed (scBasset, SIMBA) and sequence-ignorant (PeakVI and iterative LSI) methods, with and without explicit batch correction; we once again reported bootstrap 95% confidence intervals for metrics and FDR-adjusted *p*-values for pairwise comparisons (**Supplementary Fig. 4d, Supplementary Tables 2,5, Methods**). An important caveat here is that the reported labels themselves are somewhat uncertain, since the authors had to perform a difficult batch correction and clustering to annotate their dataset; embedding methods that are more similar to the approach used in the original paper may be more likely to recapitulate their results, whether the labels are correct or not. Nevertheless, for the Satpathy *et al.* large hematopoietic dataset, CellSpace significantly outperformed (adjusted *p* < 0.01) all methods except for PeakVI (batch-corrected), which outperformed CellSpace here (adjusted *p* < 0.05), even though it was one of the poorer performers on the small hematopoietic and human cortex datasets. The performance improvement was due to PeakVI’s better biological complexity score relative to reported cell type labels (adjusted *p* < 0.01); the batch correction scores for CellSpace were higher than PeakVI on average, but not significantly different.

For the Satpathy *et al.* TME dataset, to reduce potential label uncertainty, we restricted the evaluation of clustering and batch correction metrics to the non-tumor cells, excluding clusters labeled ‘tumor1’ through ‘tumor4’, although all cells were embedded by all methods (**Methods**). We found that CellSpace significantly outperformed all other methods based on batch score (adjusted *p* < 0.01 in all cases), but only outperformed batch-corrected SIMBA on biological complexity score (adjusted *p* < 0.01), with comparison to other methods giving ties or losses for this score; on overall score, CellSpace mainly gave statistical ties to other methods, with significant wins over Harmony-corrected itLSI (adjusted p < 0.01), batch-corrected SIMBA (adjusted *p* < 0.01), and PeakVI (adjusted *p* < 0.05), but a loss to batch-corrected PeakVI (adjusted *p* < 0.05) (**Supplementary Fig. 4d, Supplementary Tables 2,6**). We note however that PeakVI does not provide TF motif scores, and no other sequence-informed method (i.e., with the potential to compute batch-corrected single-cell motif scores) outperforms CellSpace.

To demonstrate scalability up to another order of magnitude in number of cells, we applied CellSpace to a very large, diverse, and multi-donor human fetal scATAC-seq atlas^20^, consisting of ∼720K cells from 20 donors in 3 batches. We used a latent space of dimension 70 to accommodate the diversity of cell types, and we computed variable peaks on a sample of the dataset (∼5% of cells) and used these events for the full-scale embedding (**Methods**). Due to our memory-efficient implementation, there were no problems training the CellSpace model (**Supplementary Table 1**). Qualitative visualization of all cells with UMAP showed meaningful relationships between more closely related tissues (**Fig. 4c**), and batches were well mixed. Moreover, blood cell types from multiple organs clustered together; for example, a cluster that prominently features lymphocytes from the thymus also includes cells labeled “lymphoid/myeloid” from placenta (**Fig. 4c**).

Finally, we applied CellSpace to combine two distinct chromatin accessibility datasets using different assays to profile the human cortex: the scATAC-seq readout of the multiome dataset presented above (**Fig. 3a**) and a scATAC-seq dataset from the same study^18^. These two datasets were processed independently to generate different peak atlases. Selecting the 50K most variable peaks in each dataset yielded both overlapping and dataset specific peaks, as depicted in the Venn diagram (**Fig. 4d**). Here, the intersection of 31,800 peaks indicates peaks with non-zero overlap, but not necessarily the same boundaries, in the two atlases. Without reprocessing these datasets from scratch to generate a combined cell-by-peak matrix relative to a common peak atlas, this situation would yield an “uncorrectable” batch effect for standard methods, which require a shared peak or tile feature representation. We trained a CellSpace embedding to combine the two datasets, each represented with respect to its own peak atlas and associated with a batch covariate—which we used to avoid pushing cells from different batches away from each other in sampling negative cells (**Methods**)—and found that batch effects were indeed corrected (**Fig. 4d**). Annotating the combined embedding using cell type annotations from the two datasets recovered the correct overall structure (**Fig. 4d**), with inhibitory and glutamatergic neurons well separated and progenitor populations like OPC and RG placed at the apex of the developmental manifold. Clustering on the CellSpace embedding identified coherent clusters that mixed cells of similar types from the multiome and scATAC-seq datasets (**Supplementary Fig. 4e,f**). This example shows the unique and powerful ability of CellSpace to integrate independently processed chromatin accessibility datasets through its sequence-informed embedding.

## Discussion

By training an embedding of both DNA *k*-mers and cells into a common latent space with a memory efficient implementation, we have shown that CellSpace learns latent structure in multi-sample and even multi-assay single cell chromatin accessibility datasets while mitigating batch effects. TF motif activities in single cells can naturally be inferred based on the similarity of TF motif and cell embeddings in the CellSpace latent space. Importantly, this motif scoring procedure does not require the TF motifs to be known at training time. In large multi-batch datasets shown here, we observed that CellSpace’s sequence-informed embedding implicitly mitigated batch effects, even without use of a batch covariate. In one case, where datasets were independently processed with respect to distinct peak atlases, we used a batch covariate simply to avoid pushing cells from separate batches away from each other when training CellSpace; this strategy allowed us to correct a batch effect that would be “uncorrectable” by any other method without reprocessing from scratch. Indeed, we have found only rare cases where a clear batch effect persists after training CellSpace. In such cases, the embedding of cells can be easily corrected by Seurat’s anchor-based data integration method^3^, inspired by the concept of mutual nearest neighbors (MNN)^21^, where CellSpace’s embedding is used to create NN graphs, identify anchors, and to correct the batch effect in this approach (**Methods**).

In our benchmarking experiments, CellSpace was overall a top performer across datasets, giving equal or significantly better performance when compared to standard LSI-based embedding methods with or without Harmony batch correction or to other sequence-based embedding methods (scBasset, SIMBA, chromVAR). Interestingly, PeakVI was one of the poorer performers on the smaller datasets, but batch-corrected PeakVI was the only method to outperform CellSpace (adjusted p < 0.05) on the larger datasets. We note that PeakVI is an embedding of the cell-by-peaks binary matrix and does not provide TF motif scores, whereas CellSpace provides TF motif scores that do not require an explicit batch correction. Importantly, no other sequence-informed method (i.e., with the potential to compute batch-corrected single-cell motif scores) outperforms CellSpace. CellSpace has impressive batch mitigation properties, with only one loss to another method in all pairwise comparisons across three datasets. While explicit batch correction (e.g., by Harmony) sometimes helps and sometimes hurts – and it is not always clear which is happening – CellSpace gives consistently strong performance without the requiring an explicit consideration of batch effects.

We have found that the default parameters—sampling 150bp sequences, generating 20 examples per event (tile or peak) per epoch, using 8-mers with 3-grams (*k* = 8, *N* = 3), embedding in a 30-dimensional latent space, and training for >20 epochs—work well in most cases. However, sometimes, experimentation with different parameters is needed to obtain a good embedding; for example, a very large and diverse dataset typically requires a higher dimensional embedding space and a larger number of epochs to train. A qualitative sign that the CellSpace representational hyperparameters need to be optimized—or possibly that longer training is needed—is a “cloudy” embedding where distinct cell types have not been pulled apart enough, as visualized in a UMAP; particularly, in the case where all cells represent the same cell type in different states, so that chromatin accessibility changes and TF motif activity differences are present but more subtle. We have found it easier to obtain a good embedding right off the bat (i.e., with minimal changes to default parameters) when using variable tiles rather than a peak atlas; possibly, the quality of the peak atlas influences the amount of parameter optimization required. Moreover, using top variable peaks or genomic tiles identified by iterative LSI, instead of a large peak atlas, significantly improves running time while preserving or possibly improving the embedding quality. We also note that when performing a Seurat SNN clustering on the CellSpace embedding, a higher resolution is often needed to obtain the same number of clusters as compared to a standard itLSI-based embedding. Additionally, while we have presented a batch-aware version of CellSpace—where negative cells are sampled within the same batch as the positive cell—as a way to integrate datasets processed with respect to different peak atlases, this batch-aware version also appears useful when using variable tiles and integrating datasets from separate studies.

While CellSpace is currently limited to scATAC-seq, we foresee an extension to multiome data where cells, genes, and *k*-mers are embedded in the same space, and cell embeddings are updated both by the current procedure of sampling sequences from peaks using the scATAC readout and also by (LHS, RHS) pairs consisting of expression-weighted gene lists and cell labels from the scRNA readout. However, use of both modalities entails weighting how much sequence features vs. gene expression features should influence the similarity of cells in the embedding space. We note that the latent semantic embedding of entities in StarSpace has also been reformulated as a graph embedding problem, where entities are vertices and (LHS, RHS) pairs specify edges in a graph^22^. This graph embedding approach has been recently proposed as the SIMBA embedding technique for scRNA-seq, scATAC-seq, and multiome data^7^; for scATAC, cells, peaks, *k*-mers, and TF motifs are all explicitly embedded as vertices, and each cell is connected by edges to its peaks. While related to our approach, CellSpace makes important algorithmic choices that are less naturally framed as a graph embedding problem. In particular, CellSpace does not explicitly embed peaks—a choice that appears to mitigate batch effects in datasets analyzed here. Indeed, SIMBA’s “peaks only” graph embedding of the cell-by-peak matrix is not so different from standard dimensionality reduction, which can suffer from technical batch effects, and in our experiments, we found that embedding of the *k*-mer and motif nodes did not eliminate these effects based on quantitative batch correction and clustering metrics. Other important CellSpace implementation choices are the sampling of negative examples, which addresses the label asymmetry in scATAC-seq (there are orders of magnitude more inaccessible events than accessible events for each cell, and some of the 0s in the cell-by-peak matrix are false negatives due to limited capture efficiency); the use of *N*-grams to capture local sequence context; and the sampling of sequences from chromatin accessible events to improve robustness. Finally, CellSpace enables the embedding of DNA sequences and TF motifs that were not explicitly introduced to the model at training time, based on the learned embedding vectors for *k*-mers, and importantly does not rely on any a priori choice of motifs.

There is also a connection between CellSpace and the recent neural network-based scBasset approach. To make this link, we can view CellSpace as implicitly embedding peak (sub)sequences to a latent space while representing every cell as a classification model vector that predicts whether the embedded sequences are accessible in that cell. In CellSpace, the classification rule for each cell is simply based on the cosine similarity between the embedded sequence and the cell model vector in the latent space. This view is made explicit in scBasset, which learns a neural network embedding of peak sequences together with cell-specific model vectors in the latent space and minimizes classification loss using the entire (binarized) cell-by-peak matrix as output labels. The neural network sequence embedding is more expressive than our *N*-gram of *k*-mers representation but may also be more prone to overfitting and learning batch-specific technical artifacts (which scBasset then explicitly attempts to model). Additionally, scBasset requires high-memory GPUs to train the neural network model in a practical running time. Moreover, there is an asymmetric label noise in the binary cell-by-peak matrix—positive labels (1s) almost certainly represent accessible peaks in individual cells, but negative labels (0s) could either be truly inaccessible or accessible but not captured in the library. By sampling from negative examples in training, CellSpace may be less susceptible to false negative events than a multitask classification approach. Still, these new sequence-informed embedding methods—CellSpace, graph embedding, and neural network multitask learning—potentially have complementary strengths that could be combined in future algorithmic innovations for discovery of latent structure in single-cell epigenomic data.

## Data Availability

The following public datasets, available through the Gene Expression Omnibus, were used in this study: GSE96769, GSE74310, GSE125523, GSE162170, GSE129785, and GSE149683.

## Code availability

CellSpace is freely available on GitHub at https://github.com/zakieh-tayyebi/CellSpace.

## Acknowledgements

This work was supported by NIH/NHGRI U01 award HG009395, NIH T32 GM132083, and by an award from the Geoffrey Beene Cancer Research Center.

## Author Contributions

ZT and CL formulated the problem. ZT developed the CellSpace algorithm and created the C++ and R libraries for CellSpace. ZT and AP trained CellSpace, ArchR, chromVAR, scBasset, SIMBA, and PeakVI embeddings. ZT performed the biological conservation and batch correction benchmarking. AP performed the TF motif score comparisons. CL supervised the research and drafted the manuscript, with all authors contributing to the text and figures.

---

**Supplementary Table 1.** Run time and memory usage information for all datasets.

**Supplementary Table 2.** Biological conservation, batch correction, and overall scores.

**Supplementary Tables 3-6.** Result of pairwise comparisons of performance metrics between all methods over 1000 bootstrap samples, including FDR-adjusted *p*-values, for **(3)** Buenrostro et al. dataset, **(4)** Trevino et al. dataset, **(5)** Satpathy et al. hematopoietic dataset, and **(6)** Satpathy et al. TME dataset.

## Methods

### CellSpace algorithm

CellSpace uses the StarSpace (mode = 0) algorithm to learn a co-embedding of DNA *k*-mers (*k* = 8 by default) and cells into a latent vector space ℝ^*d*^ (*d* = 30 dimensions by default) based on training example sequences sampled from accessible events.

Accessible events are either an atlas of accessible peaks or variable tiles, for which a cell-by-event matrix of accessibility is available. Top variable tiles (500 bp genomic bins) can be identified using ArchR’s iterative LSI method. When stated, we used top variable peaks, instead of the entire peak atlas, which were identified with an adaptation of ArchR functions.

Starting from a binary cell-by-event matrix, CellSpace creates multiple training examples per event (20 by default) while training during each epoch (50 epochs by default). To generate a training example for an event, an *L*-length (*L* = 150 bp by default) DNA sequence is randomly sampled from the corresponding genomic region. The bag of *L* − *k* + 1 consecutive overlapping *k*-mers, created by sliding a window of size *k* across the sampled sequence by 1 nucleotide at a time, is used as the ‘input’. Assuming each DNA *k*-mer and its reverse complement have identical genetic information, we hash them to the same row of the embedding matrix. The cells for which the event is accessible are used as ‘positive labels’. The model is optimized so that the ‘input’ sequence is embedded closer to its ‘positive labels’ in the latent space than to ‘negative labels’ (i.e., *K* randomly sampled cells for which the event is not accessible) which are selected by *K*-negative sampling.

StarSpace represents features, which are embedded directly, and entities (i.e., bag of one or more features) by a *d*-dimensional vector. The inferred embedding of an entity comprised of *M* features is given by 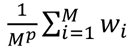 are the vector representations of its features, and *p* = 0.5 is the default value. CellSpace embeds cells (as ‘labels’) and *k*-mers (as features in ‘input’) directly and infers the embedding of any DNA sequence as a bag of *k*-mers, enabling the comparison of sequences and cells in the same space.

Additionally, CellSpace learns contextual information from the relative position of the *k*-mers by training StarSpace with *N*-grams (window of *N* = 3 consecutive *k*-mers by default), so that each pair of *k*-mers within an *N*-gram is also considered as a feature, embedded directly with a row in the embedding matrix, and added to the ‘input’ of the training example. For *N* > 1, StarSpace uses a hashing trick to retrieve the embedding vector of an *N*-gram. The user can control the size of the hashing map ‘bucket’.

At step *i* of stochastic gradient descent (SGD) optimization, StarSpace picks one random ‘positive label’ as the right-hand side entity *RHS_i_* of the training example and uses the ‘input’ as the left-hand side entity *LHS_i_* . Analogously, CellSpace randomly selects a positive cell for the corresponding event as the *RHS_i_*. The ‘input’ *L*-length sampled sequence represents the *LHS_i_*, and its embedding is inferred from the embedding vectors of its features as described above. CellSpace then samples *K* random ‘negative’ cells *c_n_* … *c_nk_*—for which the event is not accessible—and optimizes the parameters to pull the *LHS_i_* closer to the embedding of the positive cell and away from that of the negative cells by minimizing the margin ranking loss:

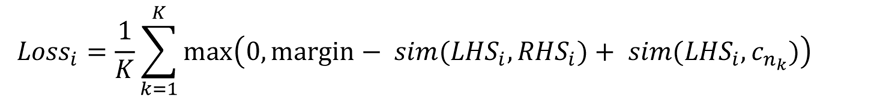

Here *sim* is the cosine similarity in the embedding space by default. Therefore, the loss increases unless the event is closer to the positive cell than the negative cell, and the difference is greater than the margin. The embedding of a negative cell is not updated if it yields zero loss, because it is already sufficiently distant to the event.

CellSpace has been integrated into the C++ StarSpace implementation so that the sparse cell-by-event matrix and the DNA sequences of the events are loaded, parsed, indexed, and stored in memory. Training example batches are randomly created in real time during training and are only temporarily stored, so that the running time of CellSpace will increase linearly with the number of training examples and the memory usage is constant. Furthermore, CellSpace utilizes the parallel training capability of StarSpace, which enables scalability to larger single-cell ATAC-seq datasets.

Multiple scATAC-seq datasets represented by different sets of events (i.e., peak/tile sets) can be simultaneously embedded by CellSpace. All datasets are initially loaded, and training examples are created in random order. The event, the positive cell, and the negative cells for each training example are sampled from the same dataset. This co-embedding utilizes the shared DNA sequence information between events that may not have the exact same genomic region.

### CellSpace visualization, clustering, and motif embeddings

CellSpace outputs embedding vectors for cells and *k*-mers after training a StarSpace model on scATAC-seq data.

The CellSpace embedding of each transcription factor (TF) motif is computed by creating a bag of *k*-mers by sliding a *k* bp window across the consensus motif sequence, and computing its embedding from the embedding vectors of its *length(motif)* − *k* + 1 constituent *k*-mers as previously described for a StarSpace entity. Cell-by-TF similarities (i.e., cosine similarity between CellSpace embedding vectors) are computed, and *z*-scored across all cells per TF, to represent TF activities.

The pairwise distance matrix of cells (i.e., cosine distance between CellSpace embedding vectors) is used to build a NN and shared nearest neighbor (SNN) graph. Cells are visualized with a UMAP embedding and clustered using the Louvain method on the SNN graph by Seurat^23^.

To visualize cells and TFs in the same space, the embedding vectors of selected TFs are concatenated to the embedding vectors of cells, and their pairwise cosine distances are used to compute a UMAP embedding as described above.

The sequence-informed embedding of CellSpace captures the structure of scATAC-seq data across multiple samples, donors, and datasets while mitigating possible batch effects. However, if a batch effect persists in the CellSpace embedding, we found the problem could be easily corrected by Seurat’s data integration method^3^. CellSpace can place multiple datasets in a shared low-dimensional space, which can be used instead of canonical correlation analysis (CCA) to identify and score pairs of MNN ‘anchors’ between datasets. Similarly, the NN graphs used for weighting the anchors for cells within each dataset can be created from the CellSpace embedding, instead of using PCA dimensionality reduction. Finally, the batch effect can be removed by correcting the CellSpace embedding of ‘query’ datasets with respect to the ‘reference’ dataset, similar to how gene expression matrices are corrected by Seurat v3.

### Discovering de novo motifs with CellSpace

We computed the inferred embedding of all possible DNA 10-mers by sliding an 8 bp window across each 10 bp sequence and computing the average CellSpace embedding of its 3 constituent 8-mers. We built a bipartite *K*=50 nearest neighbor graph between cells and 10-mers, based on their cosine distance in the embedding space, representing each 10-mer and its reverse complement as a single vertex in the graph. For each group of cells, we identified the 10-mers that were among the nearest neighbors of at least 20% of its cells. These 10-mers were clustered by kmer::cluster in R^24^, using a top-down tree-building approach, and cutting the tree at height=0.5. For each cluster of size >3, we aligned the 10-mers by msa::msaClustalW in R with default settings^25^. From each alignment, we computed the position weight matrix (PWM) of a *de novo* motif. The embedding of each *de novo* motif was computed as the average embedding of the 10-mers in its corresponding cluster.

### Evaluating scATAC-seq analysis results

#### Clustering and visualization

For each embedding, cells were clustered using Seurat’s SNN method and visualized by UMAP, with *K*=20 by default and metric=’cosine’ for CellSpace and metric=’euclidean’ for other methods. We used a range of values as Louvain clustering resolution and picked the value that yielded the same number of clusters as cell types (i.e., the cell labels that would be used as ground truth in evaluation). In a few cases where no such value was found and there were too many clusters, we merged the smallest clusters into the nearest larger clusters based on their connectivity in the SNN, using the R function CellSpace::merge_small_clusters which was adapted from Seurat::GroupSingletons.

#### Biological conservation scores

To evaluate the embedding and clustering results from each method, we used the implementation of Average Silhouette Width (ASW), Normalized Mutual Information (NMI), and Adjusted Rand Index (ARI) by scib^14^ in python, as well as the implementation of Homogeneity (HG) by sklearn^26^ in python. The biological conservation score was computed as the average of all 4 metrics.

#### Batch correction scores

To evaluate the batch effect in the embedding of each method, we used batch ASW (bASW), Graph Connectivity (GC), and *k*-nearest neighbor Batch-Effect Test (kBET) from scib. To speed-up the bootstrapping process for the large-scale hematopoietic and tumor microenvironment datasets, we used the implementation of kBET by scib-metrics in python, which approximates the method used in the original scib package and utilizes GPUs. The metric batch NMI (bNMI) was computed as 1-NMI(cluster, batch) in each cell type, and reported as the average over all cell types. The batch correction score was computed as the average of all 4 metrics.

#### Overall score

The overall score is the weighted average of the biological conservation and batch correction scores, with 0.6 and 0.4 as their relative weights, respectively.

#### Bootstrapping

For each dataset, we created *B* = 1000 bootstrap samples from the original dataset by resampling the same number of cells, with replacement. For each embedding, we clustered every bootstrap sample and computed the corresponding benchmarking scores as described above. For confidence level 1 − *α* of a statistic, we reported the percentile confidence interval, i.e., the ^*α*^ and 1 − quantiles of the bootstrap distribution. To compare the scores of two methods, we performed a two-sided test under the null hypothesis *θ* = 0, where *θ* is the difference in scores. We computed the *p*-value of the null hypothesis using confidence interval inversion: The *p*-value for a two-sided test of the 1 point-null hypothesis *θ* = *θ*_0_ is the smallest *α* ∈ [ , 1] such that *θ*_0_ is not contained in the 1 − *α* *α* confidence interval from the bootstrap distribution of *θ*. For each dataset, we performed pairwise tests between all the methods and FDR-adjusted the *p*-values.

### Additional method parameters

#### scBasset

scBasset was trained with its default Basenji-inspired architecture and a bottleneck layer size of 32^6^. For batch-correction, batch labels were provided as input to the scBasset-BC architecture, which adds a fully connected layer to predict the batch-specific contribution prior to the final sigmoid.

#### SIMBA

For the peak-only version, SIMBA was run on peak-by-cell matrices using default settings. Unless stated otherwise, the embedding was trained on peaks associated with top PCs. For the sequence-aware version, the peak set was annotated with *k*-mers and motifs using the scan_for_kmers_motifs R function, and peak-motif and peak-kmer edges were included in graph generation. To obtain motif scores, we used the compare entities function between cell embedding and motif embedding matrices, followed by subsequent softmax transformation^7^. For batch-corrected SIMBA, peak-by-cell matrices were split by batch and edges between batches were inferred using their mutual nearest neighbor implementation in the infer_edges function, and edges between batches were included in graph generation. For all versions, the model was trained for the recommended 10 epochs, at which point the validation loss leveled and the embedding had converged.

#### PeakVI

PeakVI was run with default settings (2 encoder layers, 2 decoder layers, dropout rate 0.1) on the peak-by-cell matrix as input, and optionally providing donor annotations for explicit batch correction^15^.

#### chromVAR

chromVAR ‘deviations’ of JASPAR 2020 motifs, for the motif version, or that of DNA 8-mers, for the *k*-mer version, were computed for the peak-by-cell count matrix, following standard steps with default parameters. Highly correlated features (cor>0.9) and features with low variance (sd<1.5) were removed from the cell-by-motif/kmer deviation *z*-score matrix, and principal component analysis (PCA) was performed on the filtered matrix. The PCs were used as the chromVAR embedding^4^.

### Small-scale human hematopoiesis dataset

Paired-end FASTQ files for 2,779 samples from GSE96769 and 192 LMPP/Mono samples from GSE74310^5^ were downloaded using SRA toolkit v2.11. Adapters were detected and trimmed by TrimGalore v0.6.6 (FastQC and Cutadapt wrapper). Reads were aligned to hg19 by Bowtie2 v2.4.2 with ‘’ --maxins 2000 --very-sensitive --no-unal --no-mixed”. Barcodes were added to the aligned BAM files as ‘CB’ tags using a custom script. BAM files were filtered (MAPQ>30), coordinate-sorted, indexed, and merged by Samtools v1.11.

The final BAM file was inputted to ArchR v1.0.1, barcodes were filtered (TSS enrichment score > 4 and 5K < number of fragments < 200K), and doublets were detected and removed with default settings. 2,154 cells were retained for downstream analyses. Using ArchR’s implementation of iterative LSI, the top 50K most variable tiles (genome-wide 500 bp bins) were identified after 5 iterations. This was restricted to the standard chromosomes chr1, …, chr22. The selected tiles and their corresponding cell-by-tile counts matrix were extracted for training a CellSpace model.

The result of scaling the dimensionality reduction from the last iteration of iterative LSI, named ‘ArchR itLSI (var. tiles)’, was used to build a NN graph and its SNN graph and to cluster the cells (method=“Seurat”) to identify ArchR clusters. The ArchR UMAP embedding was computed from the NN graph. The itLSI embedding was batch-corrected using Harmony^16^ with ‘Donor’ annotations as batch labels.

A CellSpace model was trained with “--k 8 --sampleLen 150 --dim 30 --ngrams 3 --exmpPerPeak 20 -- epoch 50” on the cell-by-tile count matrix. The resulting CellSpace embedding, named ‘CellSpace (var. tiles)’, was used for downstream analyses.

Pseudotime analysis was performed on the CellSpace embedding by standard functions from Palantir v1.0.0^12^. An HSC cell was manually selected as the starting point, but the potential trajectories and their terminal cells were identified by Palantir.

Moreover, we downloaded the peak atlas and cell-by-peak count matrix for the 2,154 cells, filtered as described previously, and retained 277,916 promoter-distal peaks from the standard chromosomes chr1, …, chr22 that were accessible in at least 5 cells.

The result of scaling the LSI reduced dimensions computed from the filtered count matrix was named ‘LSI (peaks)’. The LSI embedding was batch-corrected using Harmony with ‘Donor’ annotations as batch labels.

Another CellSpace embedding, named ‘CellSpace (peaks)’, was trained with “--k 8 --sampleLen 150 -- dim 30 --ngrams 3 --exmpPerPeak 20 --epoch 50” on the count matrix of the top 50K most variable peaks, which were identified by performing 5 iterations of iterative LSI. We did not correct for any possible batch effects during this process.

scBasset was run on the cell-by-peak matrix, with default settings, and the embedding was named ‘scBasset (peaks)’. Peaks accessible in fewer than 5% of cells were removed per their default filtering. The embedding ‘scBasset (batch-corrected)’ was trained as described previously, with ‘Donor’ annotations as batch labels.

We performed different versions of SIMBA on the filtered count matrix. The peak-only embedding and the sequence-aware embedding were named ‘SIMBA (peaks)’ and ‘SIMBA (peaks+kmers+motifs)’, respectively. Furthermore, we computed a SIMBA embedding, named ‘SIMBA (batch-corrected)’, for which ‘Donor’ annotations were explicitly encoded as batch labels.

The PeakVI embedding, named ‘PeakVI (peaks)’, and the PeakVI embedding corrected for the ‘Donor’ batch effect, named ‘PeakVI (batch-corrected)’, were computed from the filtered count matrix.

We computed a chromVAR embedding using motifs, named ‘chromVAR (peaks+motifs)’, and one using DNA *k*-mers, named ‘chromVAR (peaks+*k*mers)’.

We compared the results, excluding the “unknown” cell type, as described in ‘*Evaluating scATAC-seq analysis results*’ (**Supplementary Fig. 2d,e** and **Supplementary Tables 2,3**). ’Cell type’ labels were used as ground truth to evaluate clustering results, and ‘Donor’ labels were used as batch labels (**Supplementary Fig. 2a-c**).

### Mouse mammary epithelial dataset

The promoter-distal peaks and the cell-by-peak raw count matrix were downloaded from https://github.com/jaychung10010/Mammary_snATAC-seq (GSE125523)^17^. The t-SNE embedding from the original study, computed from LSI reduced dimensions, was also downloaded (**Fig. 2f**).

The peaks from the standard chromosomes chr1, …, chr19, chrX that were in the upper 90% quantile in their total number of counts were retained for training a CellSpace model.

A CellSpace model was trained with “--k 8 --sampleLen 300 --dim 30 --ngrams 3 --exmpPerPeak 20 -- epoch 20” on the 7,846 cell by 130,887 peak count matrix. We trained two additional CellSpace embeddings with “--ngrams 1” and --ngrams 5” and otherwise similar parameters.

### Human cortex multiome dataset

10X Multiome ATAC and RNA count matrices for human developing cerebral cortex sample (PCW21) were downloaded from GEO (GSE162170)^18^. The dataset consisted of 8,981 cells and 467,315 peaks with 14 identified cell types.

10X Multiome RNA data was processed with standard steps: genes expressed in fewer than 3 cells were discarded, counts were normalized to 10,000 reads per cell and then log transformed.

The LSI embedding was computed from the cell-by-peak count matrix.

A CellSpace embedding was trained with “--k 8 --sampleLen 150 --dim 30 --ngrams 3 --exmpPerPeak 10 --epoch 30”. Peaks not accessible in any cells were not included.

scBasset was trained as described previously. Peaks accessible in fewer than 5% of cells were removed per their default filtering, leaving 38,502 peaks. scBasset was trained for 45 epochs, at which point the validation loss and AUC had leveled and the correlation between the intercept and the library size was 0.99 (**Supplementary Fig. 3a**).

A peak-only embedding and a sequence-aware embedding were trained with SIMBA, as described previously.

The PeakVI embedding was trained on the count matrix with default settings. Moreover, we computed a chromVAR embedding using JASPAR 2020 motifs.

’Cell type’ labels were used as ground truth to evaluate clustering results, as described in ‘*Evaluating scATAC-seq analysis results*’ (**Supplementary Fig. 3d,e** and **Supplementary Tables 2,4**).

To evaluate the transcription factor (TF) activity scores of each method for single-cell multiome data, we correlated the motif scores computed from the ATAC-seq readout with the gene expression of the corresponding TFs from the RNA-seq readout. For this evaluation, all methods used JASPAR 2020^27^ as the motif database. Each TF may have multiple motifs in this database, so for each method, we selected the motif with the highest correlation to represent the TF.

Transcription factors important in cortical development, as identified in Figure 3F of Trevino et al.^18^, were highlighted in red (PAX6, INSM1, SOX9, EMX2, LHX2, FOXG1, TCF4, TCF3, TFAP2C, FOS, POU3F3, NEUROD2, BHLHE22, JUND, MEF2C, POU2F2, NFIA, MEIS2, EOMES). Motifs were filtered for those present in peaks that were accessible in at least 5% of cells, due to the default peak filtering by scBasset.

#### Combination of multiple human cortex datasets

The multiomic profiles of 8,981 human cortex cells, described in the previous section, as well as the scATAC-seq profiles of 31,304 human cortex cells, were downloaded from GEO (GSE162170)^18^. From the scATAC-seq dataset, we retained 12,675 cells from the time point present in the multiome dataset (post-conception week 21).

For each dataset, respectively, we identified the top 50K most variable peaks after 3 iterations of iterative LSI. The cell-by-peak count matrices of both datasets were used to train a co-embedding with CellSpace.

A CellSpace model was trained on both datasets simultaneously, with “--k 8 --sampleLen 150 --dim 50 --ngrams 3 --exmpPerPeak 20 --epoch 20”. The resulting CellSpace embedding was used for downstream analyses, without any batch correction.

#### Large-scale hematopoietic and tumor microenvironment datasets

Single-cell ATAC-seq profiles of human hematopoiesis and peripheral blood, consisting of 63,882 cells, and of human BCC TME, consisting of 37,818 cells, were downloaded from GEO (GSE129785)^19^. This includes 2,076 cells from the smaller human hematopoiesis dataset, previously described in ‘*Small-scale human hematopoiesis dataset’*.

The UMAP projection of the hematopoietic dataset from the original study, computed from LSI reduced dimensions with custom batch correction, was also downloaded (**Fig. 4a**).

For each dataset, respectively, we performed 5 iterations of iterative LSI to identify the top 50K most variable peaks. We did not correct for any possible batch effects during this process. The result of scaling the dimensionality reduction from the last iteration of iterative LSI was used as the itLSI embedding. The itLSI embedding was batch-corrected using Harmony with ‘Source’/‘Patient’ annotations as batch labels.

A CellSpace model was trained with “--k 8 --sampleLen 150 --dim 70 --ngrams 3 --exmpPerPeak 100 - -epoch 100” for each dataset, using the count matrix of top variable peaks.

scBasset was trained for 1000 epochs on the count matrix of top variable peaks, with default settings. The batch-corrected scBasset embedding was trained with ‘Source’/‘Patient’ annotations as batch labels.

A peak-only embedding, a sequence-aware embedding, and an embedding corrected for ‘Source’/’Patient’ batch effect were trained on the count matrix of top variable peaks with SIMBA. We did not restrict the training to peaks associated with top PCs.

The PeakVI embeddings with and without correction for ‘Source’/‘Patient’ batch effect were computed from the count matrix of top variable peaks as described previously.

We compared the results as described in ‘*Evaluating scATAC-seq analysis results*’ (**Supplementary Fig. 4d** and **Supplementary Tables 2,5,6**). ’Cell type’ labels (**Fig. 4b**, **Supplementary Fig. 4c**) were used as ground truth to evaluate clustering results. We used ‘Source’/‘Patient’ as batch labels (**Supplementary Fig. 4a,c**). The 4 tumor clusters were excluded from the evaluations for the TME dataset.

For 30,211 hematopoietic cells annotated as NK and T cells, we identified the top 50K most variable peaks and trained a CellSpace model with “--k 8 --sampleLen 150 --dim 30 --ngrams 3 --exmpPerPeak 50 --epoch 30”.

#### Large-scale human fetal dataset

Single-cell profiles of chromatin accessibility with three-level combinatorial indexing (sci-ATAC-seq3) from 15 human fetal organs, consisting of 720,613 cells, were downloaded from GEO (GSE149683)^20^.

To find variable peaks for this dataset, we sampled ∼5% of cells from each organ (36,200 cells total), removed peaks that were accessible in <10 cells and peaks from chrY or chrM (leaving 1,048,007 peaks), and performed 5 iterations of iterative LSI to identify the top 100K most variable peaks of the down-sampled count matrix. We did not correct for any possible batch effects during this process.

A CellSpace model was trained for all ∼720K cells and variable peaks, with “--k 8 --sampleLen 150 -- dim 70 --ngrams 3 --exmpPerPeak 50 --epoch 300”.

---

**Supplementary Figure 1.**
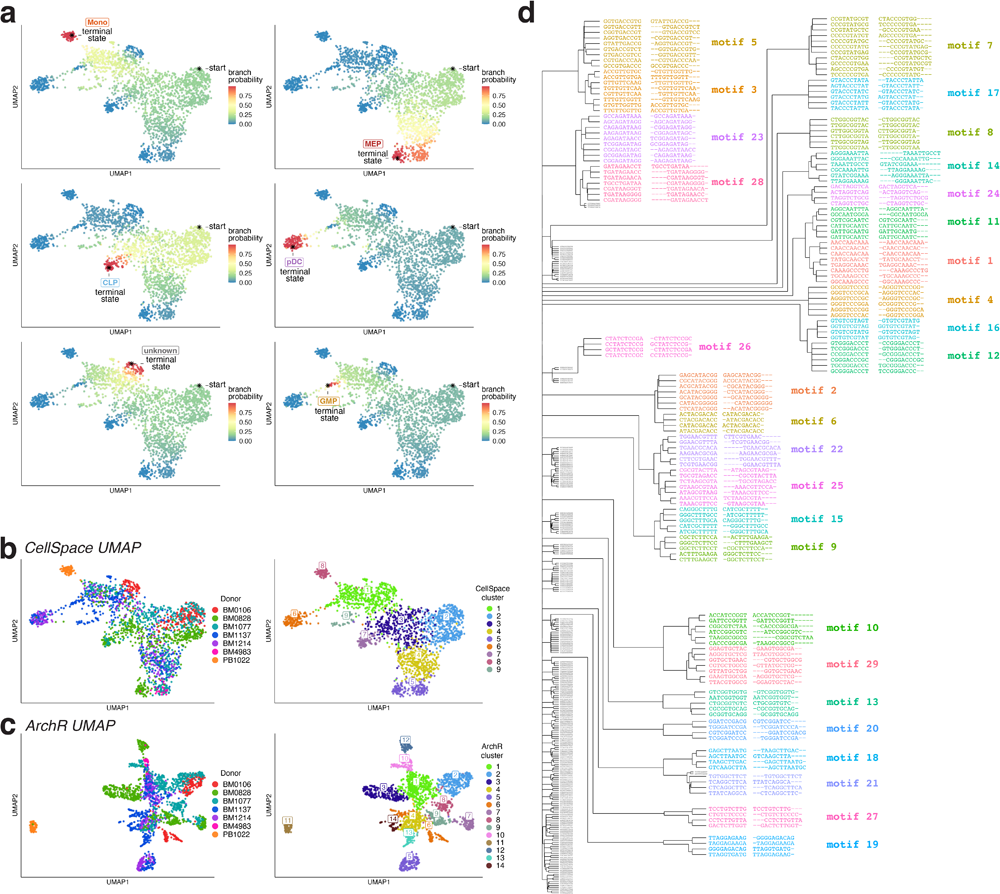
CellSpace recovers latent structure and developmental hierarchies. **a.** Palantir branch probability showing trajectories to 6 termini in the hematopoietic dataset, including termini at CLP, pDC, GMP and “unknown” GMP-adjacent population, MEP, and monocytes. **b.** CellSpace embedding annotated by donor (left) and by Seurat SNN clustering (right), which largely recovers annotated cell types. **c.** ArchR embedding of HSPC dataset annotated by donor (left) and by Seurat SNN clustering (right). **d.** Clustering of 10-mers retrieved as frequent nearest neighbors of cell clusters from the HSPC data set; 10-mers in each cluster are aligned and then converted to PWMs of *de novo* CellSpace motifs (visualized in Fig. 2e).

**Supplementary Figure 2.**
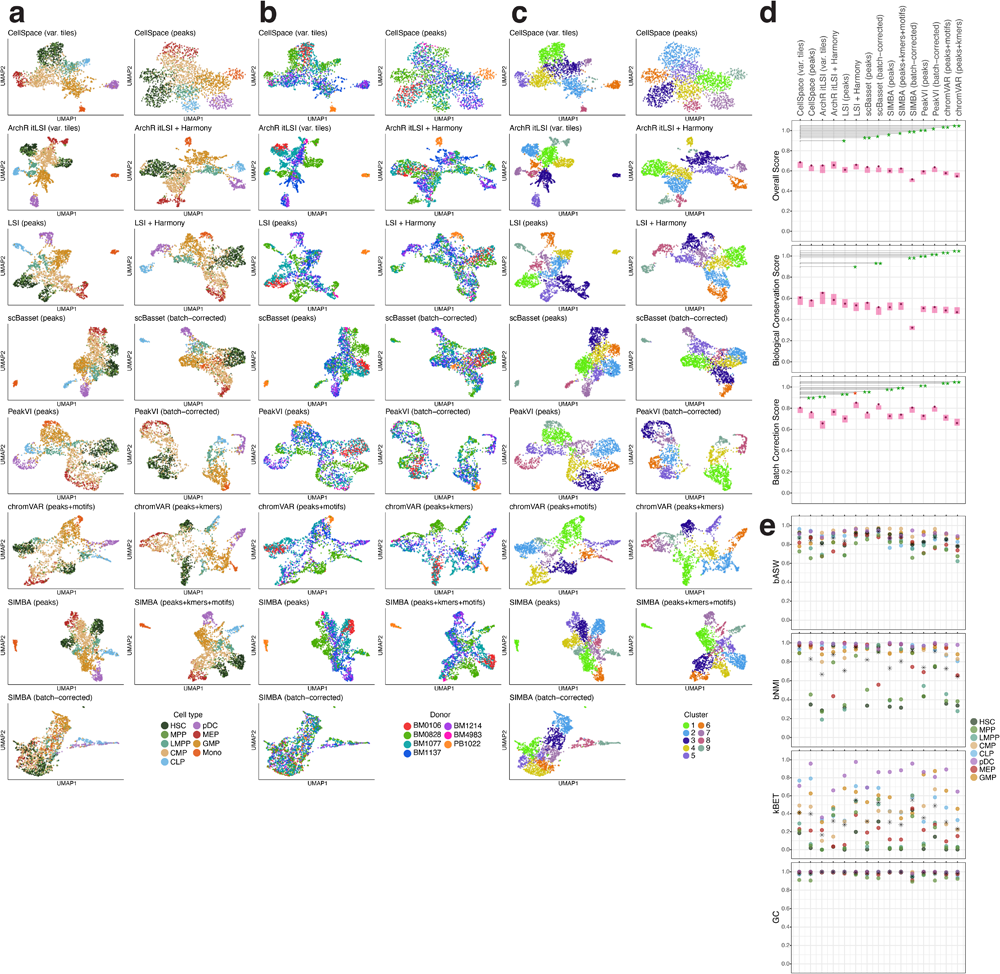
CellSpace outperforms other scATAC-seq embedding methods for batch mitigation/correction while preserving biological complexity. **a.** UMAP visualizations for multiple scATAC-seq dimensionality reduction and embedding methods on the Buenrostro et al. HSPC data set, excluding the “unknown” cell type. If the method offers an explicit batch correction option, the embedding corrected for donor batch effect is labeled as “batch-corrected”. **b.** UMAP visualizations annotated by donor (batch). **c.** UMAP visualizations annotated by Seurat SNN cluster. **d.** Performance metrics (aggregated biological conservation score, aggregated batch correction score, and overall score) for all methods on Buenrostro et al. hematopoietic data set, excluding the “unknown” cell type, with 95% confidence intervals over 1000 bootstrap samples. FDR-corrected two-sided bootstrap *p*-values for comparison against CellSpace (variable tiles) are shown. Green stars: significant win for CellSpace; red stars: significant win for other method. *: adjusted *p* < 0.05, ** adjusted *p* < 0.01. **e.** Batch correction metrics reported per cell type, excluding the monocyte cell type which consists of a single batch. Average score over all cell types is also shown.

**Supplementary Figure 3.**
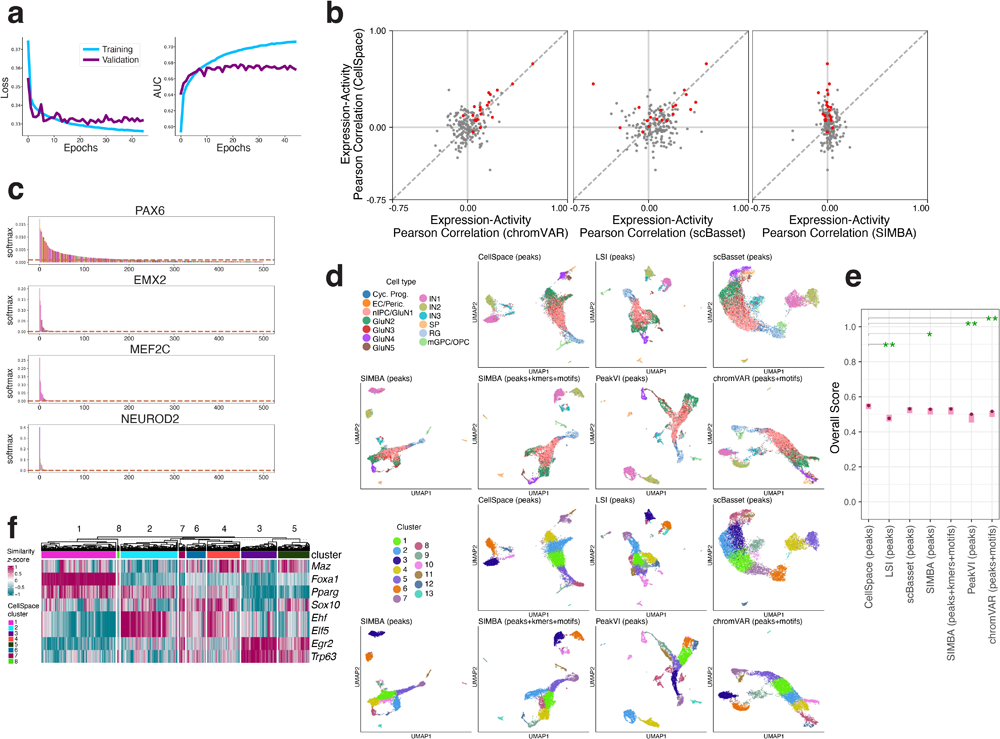
Single cell motif scoring using CellSpace accurately maps TF activities. **a.** The scBasset model converges after 40 epochs on the human cortex multiome dataset. **b.** Comparison of CellSpace vs. scBasset TF motif activity scores, CellSpace vs. SIMBA scores, and CellSpace vs. chromVAR scores based on correlation with gene expression in the human cortex multiome dataset. Important neurodevelopmental TFs shown in red. **c.** SIMBA motif scores for PAX6, EMX2, MEF2C, and NEUROD2 can be used to rank cells and learn an association with the top-ranked cell type. **d.** UMAP embedding and Seurat SNN clustering of the human cortex multiome dataset using multiple scATAC-seq embedding methods. **e.** Overall biological conservation score for all methods on Trevino et al. human cortex data set (single batch), with 95% confidence intervals over 1000 bootstrap samples. FDR-corrected two-sided bootstrap *p*-values for comparison against CellSpace are shown. Green stars: significant win for CellSpace. *: adjusted *p* < 0.05, ** adjusted *p* < 0.01. **f.** TF motif scores from the CellSpace embedding for the mammary epithelial dataset (embedding and clusters visualized in Fig. 2h).

**Supplementary Figure 4.**
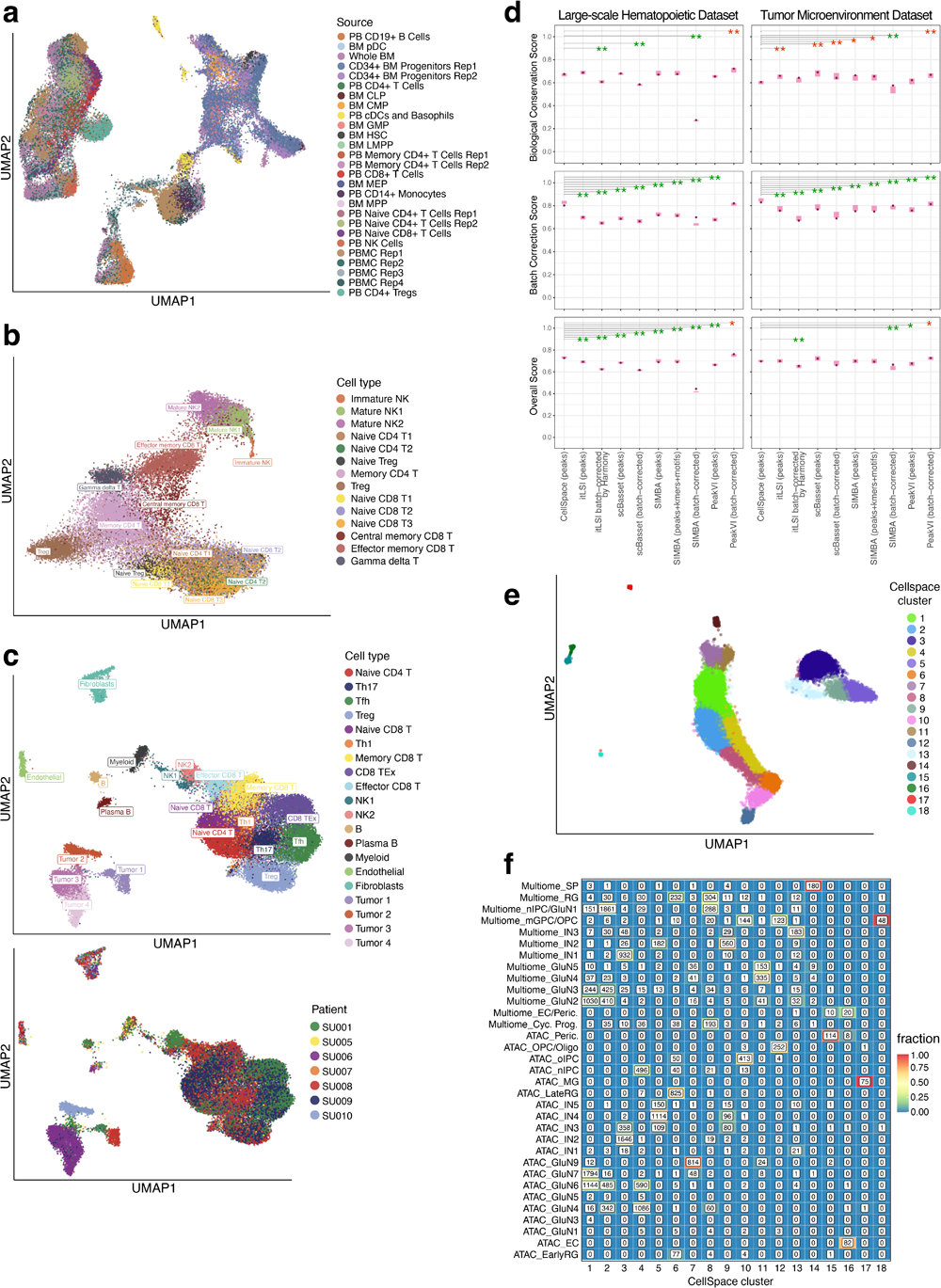
CellSpace’s embedding implicitly mitigates donor-and assay-specific batch effects in large-scale scATAC-seq datasets. **a.** Batches and human donors are well mixed in the CellSpace embedding of the large human hematopoietic dataset (visualized in Fig. 4b). **b.** CellSpace embedding of large hematopoietic dataset restricted to 30,211 NK and T cells. **c.** CellSpace embedding of 37,818 cells from BCC TME scATAC-seq data from 7 patients, annotated by cell type and by donor, recovers immune and stromal cell types with no evident donor batch effect. **d.** Performance metrics (aggregated biological conservation score, aggregated batch correction score, and overall score) for all methods on the Satpathy et al. large hematopoietic data set and the TME data set, excluding the tumor clusters, with 95% confidence intervals over 1000 bootstrap samples. FDR-corrected two-sided bootstrap *p*-values for comparison against CellSpace are shown. Green stars: significant win for CellSpace; red stars: significant win for other method. *: adjusted *p* < 0.05, ** adjusted *p* < 0.01. **e.** Seurat SNN clustering after CellSpace joint embedding of multiome and scATAC-seq human cortex datasets. **f.** Membership of annotated cell types from multiome and scATAC-seq human cortex datasets in CellSpace clusters as shown in (**e**), after joint embedding, showing coherent clusters with membership from both assays.

